# FastaGuard: FASTA preflight QC for modern bioinformatics pipelines

**DOI:** 10.64898/2026.10.28.761563

**Authors:** Ehsan Estaji, Jian-Feng Mao

## Abstract

**Summary:** FastaGuard is a streaming, explainable FASTA preflight QC tool that validates assembly FASTA files, detects structural and composition red flags, and emits a versioned multi-format report with an explicit pipeline gate before expensive downstream analysis. We evaluated v1.1.0 in four analyses whose design was fixed before data collection. On 700 public NCBI assemblies the pipeline gate stopped 19.7% (1.9% weighted to the population), always on its high-N rule and mostly in scaffold-level genomes; the NCBI submission gate stopped 2.0%, and an independent recount confirmed every stop. Among 1 040 faults injected into real assemblies, the NCBI gate flagged 520 of 520 in 13 error classes (table2asn: 280 at error level) and blocked none of 120 benign changes. Across seven downstream tools, 46 of 182 defect cells failed and 12 altered the records without a warning, all in classes the pipeline gate blocks. Stopping a mismatched reference at the gate avoided about 54–62 s of mapping and sorting at a cost of 0.05–0.06 s.

**Availability and Implementation:** Source code and documentation are freely available under the MIT license at https://github.com/ehsanestaji/FastaGuard (release tag v1.1.0: https://github.com/ehsanestaji/FastaGuard/releases/tag/v1. 1.0; archived at doi:10.5281/zenodo.23728753). Install via Bioconda (mamba install -c conda-forge -c bioconda fastaguard=1.1.0), BioContainers, or GitHub release binaries. Implemented in Rust; supported on Linux and macOS, including Bioconda builds for linux-64, linux-aarch64, osx-64, and osx-arm64.

**Supplementary information:** Supplementary Note (designs, all results, deviations from the pre-registered protocol); scripts, manifests and result tables at Zenodo, doi:10.5281/zenodo.23729530.

## 1 Introduction

Genome assemblies and reference FASTA files are foundational inputs for bioinformatics pipelines, yet FASTA-level checks are usually assembled from separate utilities. Teams commonly combine ad hoc scripts, general sequence utilities such as SeqKit (Shen et al., 2016, 2024), assembly evaluators such as QUAST (Gurevich et al., 2013), biological completeness tools such as BUSCO (Simão et al., 2015), genome-quality estimation with CheckM (Parks et al., 2015), contamination-oriented exploration with BlobToolKit (Challis et al., 2020), and late-stage report aggregation with MultiQC (Ewels et al., 2016). Raw-read QC tools remain essential before assembly, but they do not answer whether a finished assembly FASTA is structurally sound for the next stage. Each of the tools above is valuable for its intended question; none is designed as a first-step gate for a narrower but frequent need: whether a FASTA file is structurally valid, alphabet-sane, interpretable, and safe to pass into expensive downstream steps.

The cost of skipping that earlier layer is practical. Malformed headers, empty records, duplicate identifiers, invalid IUPAC characters, extreme ambiguity, and high-N scaffolds may be caught only when a downstream tool reaches the offending record, or be silently normalised along the way; we measure how often that happens for seven tools below. Pipeline authors then reimplement the same checks, with incompatible report shapes and unclear stop/go semantics for workflow engines such as Nextflow (Di Tommaso et al., 2017) and Snakemake (Mölder et al., 2021). Dedicated validators exist: py_fasta_validator (Edwards, 2023) and fa-lint (GallVp, 2025) check headers, alphabet, and identifier uniqueness but, to our knowledge, provide no structured per-finding report, and AssemblyQC embeds such input checks inside a single pipeline (Rashid et al., 2024). Statistics toolkits such as SeqKit, SeqFu (Telatin et al., 2021), and gfastats (Formenti et al., 2022) mainly summarise sequences and define no gate semantics. To our knowledge, none combines severity-graded findings, an explicit blocking-versus-advisory gate, and a schema-versioned machine-readable report (JSON, TSV, MultiQC) that workflow engines can branch on.

FastaGuard addresses that gap as a dedicated *preflight* layer (Fig. 1). It is intentionally not a competitor to interpretive assembly QC: it does not claim biological completeness, contamination taxonomy, or repository acceptance, and it does not replace official NCBI, ENA, or DDBJ validators. Instead it answers whether the FASTA itself is ready for those tools, and it returns a versioned PASS/WARN/FAIL contract, with verdicts, gates, and findings that are deterministic for a given input and options, that humans and Nextflow, Snakemake, or nf-core (Ewels et al., 2020) workflows can consume without scraping logs or HTML. The design is therefore a contract as much as a report: field names, documented finding identifiers, provenance checksums, and gate continuation booleans are intended to remain stable within a schema major version. We describe the tool and then evaluate it on public assemblies, on faults injected into real assemblies, against seven downstream tools and in a workflow gate-placement experiment.

**Figure 1.**
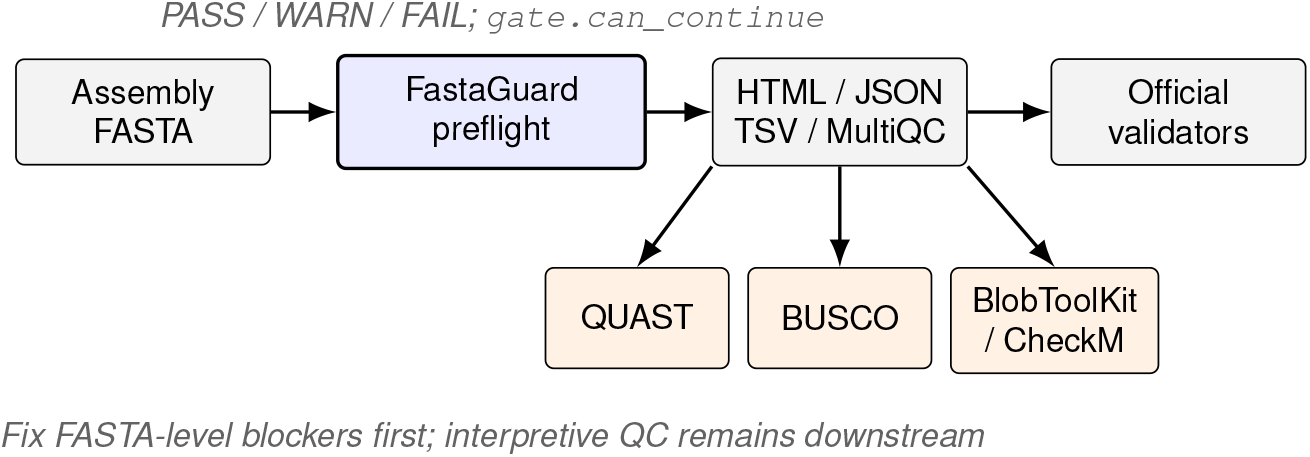
FastaGuard as the FASTA preflight layer before interpretive assembly QC and official submission validation. Machine-readable reports support workflow stop/go decisions without replacing QUAST, BUSCO, BlobToolKit, CheckM, MultiQC, or repository validators.

## 2 Implementation

FastaGuard is a streaming-first command-line tool implemented in Rust and distributed as a single binary. The primary assembly workflow is:

~~~
fastaguard sample.fa --profile assembly --gate pipeline \
   --outdir reports --prefix sample-01
~~~

which writes a four-report bundle (`.html`, `.json`, `.tsv`, and MultiQC-compatible `_mqc.json`) under a chosen prefix. Verdicts, gates, and findings are deterministic for a given input and options; provenance timestamps and durations vary between runs. JSON is the source of truth for machine routing. A committed JSON Schema and finding catalog document field names, finding identifiers, severity semantics, and suggested actions (`fastaguard --schema`, `--finding-catalog`, `--explain-finding`).

### Validity and composition checks

Under the assembly profile, FastaGuard reports structural failures (malformed headers, empty records, sequence before the first header, empty input), duplicate identifiers and duplicate sequences, invalid nucleotide/IUPAC symbols, core length statistics including N50/N90 and L50/L90, and composition signals (GC/AT/N and ambiguity rates, gap runs, tiny-contig advisories, and composition or length outlier hints). Findings are explainable: each carries an identifier, evidence bounds, and routing language that distinguishes blockers from advisories.

### Pipeline gates and process exit contract

Optional gates make stop/go policy explicit. The recommended workflow preset `--gate pipeline` marks duplicate identifiers (full-header or first-token), invalid characters, invalid FASTA structure, and high-N content as blocking by default, while GC and length outliers remain advisory routing signals unless elevated with `--fail-on`. JSON exposes `gate.status`, `gate.blocking_findings`, `gate.advisory_findings`, `gate.can_continue`, and `provenance.input_sha256`. From v0.6 onward, process exit codes separate execution from QC status: exit 0 means report generation completed for PASS, WARN, and FAIL results; exit 2 covers argument errors; exit 3 covers configuration, I/O, or runtime failures. Workflows should therefore branch on `gate.can_continue` (or related gate fields), not on process status alone. `machine_summary.safe_for_downstream` is intentionally stricter (true only for overall PASS) and is not a synonym for gate continuation.

### Submission-readiness, compare mode, and reference checks

`--gate submission` with `--submission-target generic|ncbi` performs FASTA-level submission preflight before official validators. For NCBI genome targeting, FastaGuard applies a documented, deliberately conservative `ncbi_genome` policy snapshot (first-token SeqID syntax and length limits, minimum record length, terminal Ns) based on NCBI submission and FASTA guidance (National Center for Biotechnology Information, 2025, 2026a); in a local differential check it was stricter than table2asn (National Center for Biotechnology Information, 2026b) on several boundary cases (for example, a 199-bp record or a 50-character SeqID), and it does not replace NCBI FCS (Astashyn et al., 2024), annotation validation, or repository acceptance decisions. `fastaguard compare` triages many FASTA files into cohort HTML/JSON/TSV outputs for starter cohort routing before interpretive QC. A separate `reference` mode checks whether explicitly supplied companion declarations (FASTA index, sequence dictionary, alignment/variant headers, GFF3 sequence declarations) agree with a canonical FASTA under `strict`, `coordinate`, or `advisory` policies.

### Readiness categories

Beyond a single verdict, reports expose readiness facets spanning file, structure, alphabet, index, assembly, submission, and machine concerns. That decomposition helps operators see *why* a file is not yet ready (for example, identifier hazards versus composition advisories) without reading an unstructured log. HTML reports remain human-oriented, while JSON/TSV remain stable for engines and for MultiQC custom content.

### Packaging and workflow adoption

FastaGuard is published on Bioconda (Grüning et al., 2018) and BioContainers (da Veiga Leprevost et al., 2017), ships GitHub release binaries for x86_64 Linux and for Intel and Apple-silicon macOS (Linux aarch64 is available through Bioconda), and has merged nf-core and Snakemake wrapper integrations for assembly preflight. A typical install path is `mamba install -c conda-forge -c bioconda fastaguard=1.1.0` (matching GitHub release tag `v1.1.0`), with a pinned BioContainers image for containerized workflow engines. These distribution paths are part of the Application Note claim: the tool is meant to be installed and gated in pipelines with low friction, not only built from source by specialists; we make no claim about third-party adoption. The v1.1.0 release is archived on Zenodo and citable (Estaji and Mao, 2026) (all versions: doi:10.5281/zenodo.23622310), and source and published packages remain openly reachable under the MIT license.

## Results

### Contract demonstrations

Committed public-evidence summaries for FastaGuard v0.6.0 illustrate the intended behaviour on synthetic contract fixtures and two exact NCBI RefSeq assemblies (O’Leary et al., 2016) (Table 1). Portable JSON and TSV summaries in the repository record input byte sizes, SHA-256 digests, structural metrics, finding identifiers, and scoped provenance separately for the observed executable and the verified source tree. On a recorded macOS arm64 run, a clean *Escherichia coli* K-12 MG1655 (Blattner et al., 1997) assembly (`GCF_000005845.2`; 4 641 652 bp) completed pipeline-gated preflight in 0.0569 s with PASS and no findings. A *Neurospora crassa* OR74A (Galagan et al., 2003) assembly (`GCF_000182925.2`; 41 102 378 bp) completed in 0.3441 s with WARN for gap-pattern advisories and `gate.can_continue` still true under the pipeline gate, routing to deeper assembly or composition follow-up rather than hard failure. An intentionally broken local fixture returned FAIL with blocking findings (duplicate identifiers, invalid characters, high-N content) while still exiting 0 after writing a complete FAIL report—matching the v0.6 workflow contract that separates QC status from process success. Re-running all five evidence-pack inputs with a build of the v1.0.0 tag reproduced the same verdicts, gate statuses, blocking findings, lengths, sequence counts, input SHA-256 digests, and exit status; timings differed between runs and are indicative only.

**Table 1.**
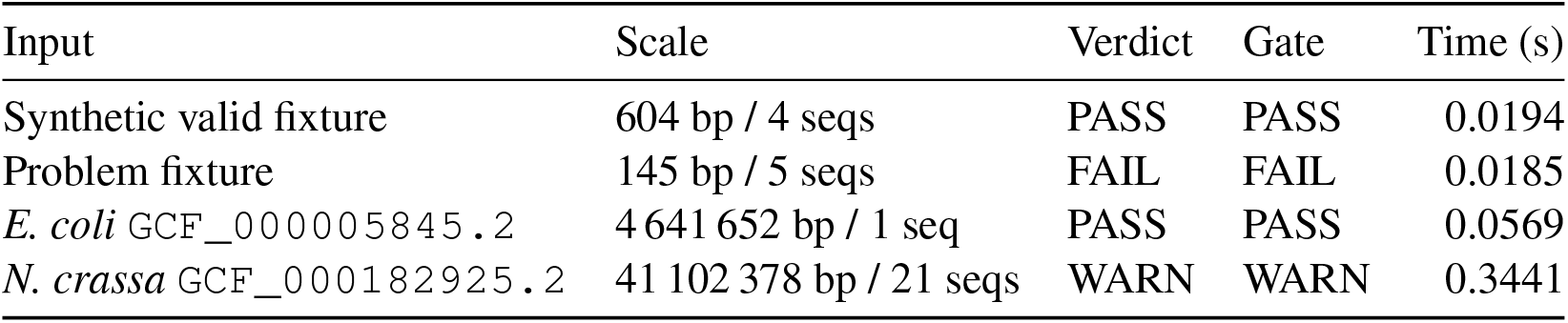
Frozen FastaGuard v0.6.0 public-evidence outcomes (assembly profile, pipeline gate) from docs/evidence/results/v0.6/evidence_summary.tsv. All runs used `--min-contig-length 1`; with the default minimum, the 604-bp synthetic fixture returns WARN through an advisory tiny-contig finding. Elapsed times are single-machine contextual measurements (macOS arm64), not cross-platform guarantees; the evaluation below used v1.1.0. The *N. crassa* count includes the mitochondrial sequence NC_026614.1 (64 840 bp).

These examples are contract demonstrations, not a representative biological benchmark across taxa or assemblers. They show that clean references can pass quickly, that advisory gap patterns can WARN without blocking continuation, and that FASTA-level blockers remain machine visible in FAIL reports that workflow engines can parse. Compare mode and submission gates extend the same contract to cohort triage and FASTA-level submission hazards; reference mode adds companion-declaration compatibility checks when indexes, dictionaries, or headers must agree with a canonical FASTA before mapping or annotation workflows proceed.

### Evaluation on public assemblies, injected faults and downstream tools

We evaluated FastaGuard v1.1.0 in four analyses whose design, predictions and analysis plan were written down and hashed before any data were collected (Supplementary Note; every deviation is listed there). In a survey of 700 public NCBI assemblies (seven taxonomic groups; half RefSeq, half GenBank-only), the pipeline gate stopped 138 of 700 (19.7%), against our prediction of under 5%; every stop was the high-N rule, none occurred among 239 contig-level assemblies, and 109 of 247 (44.1%) of scaffold-level assemblies were stopped, so the default is a policy that many scaffolded genomes trip, not a sign of corruption. Weighted to the population of eligible assemblies the rate was 1.9%. The NCBI submission gate stopped 14 of 700 (2.0%), and recounting every stop without FastaGuard code confirmed it. Record count, length and N50 matched SeqKit on all 702/702 runs. WARN is not selective (521 of 700), so workflows should branch on `gate.can_continue`. py_fasta_validator, fa-lint and SeqKit flagged none of 401 surveyed assemblies; table2asn flagged 138 of 401 (34.4%), all for undeclared N runs. When we injected 13 classes of error into 20 real assemblies (1 040 files with ground truth), the NCBI gate flagged 520/520 (100.0%), the pipeline gate blocked 360/520 (69.2%), table2asn reported an error for 280/520 (53.8%) (and no result for 120 files), fa-lint flagged 320/520 (61.5%), and py_fasta_validator and SeqKit 200/520 (38.5%) (Fig. 2); none of 120 benign changes was flagged by any validator, or by either FastaGuard gate at its blocking level. These are capability figures on author-chosen faults, not accuracy on real submissions. Of 182 cells (13 defects, two backgrounds) in seven downstream tools (samtools (Li et al., 2009), bwa (Li and Durbin, 2009), minimap2 (Li, 2018), BLAST+ (Camacho et al., 2009), Picard (Broad Institute, 2019), QUAST (Gurevich et al., 2013) and BUSCO (Simão et al., 2015)), 46 failed and 24 altered the records, 12 of them without any warning; all 12 silent alterations were in classes the pipeline gate blocks, and CRLF was harmless to all seven. Stopping a mismatched reference at the gate cost 0.05–0.06 s and avoided 54–62 s of mapping and sorting on real reads (ten matched pairs per engine), a result limited to that runner and fixture (Table 2). An earlier 30-file, author-constructed comparison, kept in the repository, gave a similar ordering of the tools.

**Table 2.**
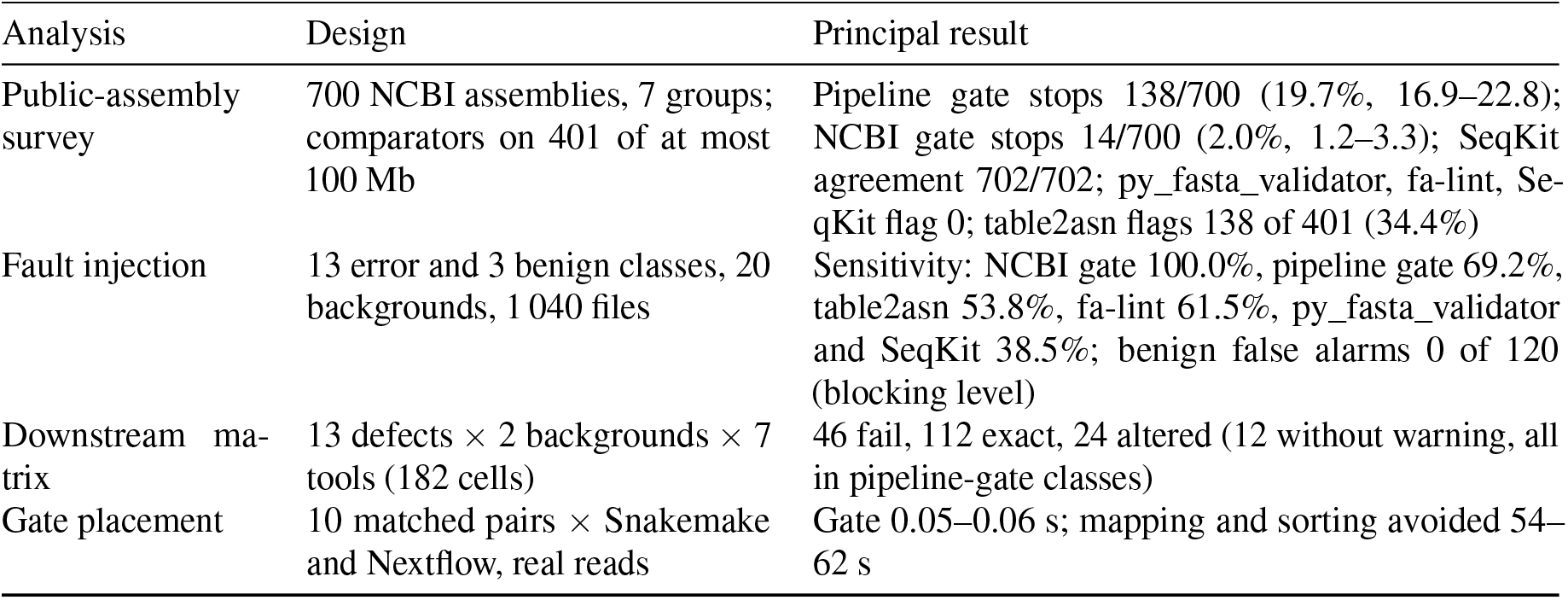
Summary of the four evaluation analyses (FastaGuard v1.1.0; details, intervals and deviations in the Supplementary Note). Intervals are 95% Wilson intervals; bootstrap intervals over backgrounds are degenerate for most tools and are given in the Supplementary Note.

| Analysis | Design | Principal result |
|---|---|---|
| Public-assembly survey | 700 NCBI assemblies, 7 groups; comparators on 401 of at most 100 Mb | Pipeline gate stops 138/700 (19.7%, 16.9–22.8); NCBI gate stops 14/700 (2.0%, 1.2–3.3); SeqKit agreement 702/702; py_fasta_validator, fa-lint, Se-qKit flag 0; table2asn flags 138 of 401 (34.4%) |
| Fault injection | 13 error and 3 benign classes, 20 backgrounds, 1 040 files | Sensitivity: NCBI gate 100.0%, pipeline gate 69.2%, table2asn 53.8%, fa-lint 61.5%, py_fasta_validator and SeqKit 38.5%; benign false alarms 0 of 120 (blocking level) |
| Downstream ma-trix | 13 defects × 2 backgrounds × 7 tools (182 cells) | 46 fail, 112 exact, 24 altered (12 without warning, all in pipeline-gate classes) |
| Gate placement | 10 matched pairs × Snakemake and Nextflow, real reads | Gate 0.05–0.06 s; mapping and sorting avoided 54–62 s |

**Figure 2.**
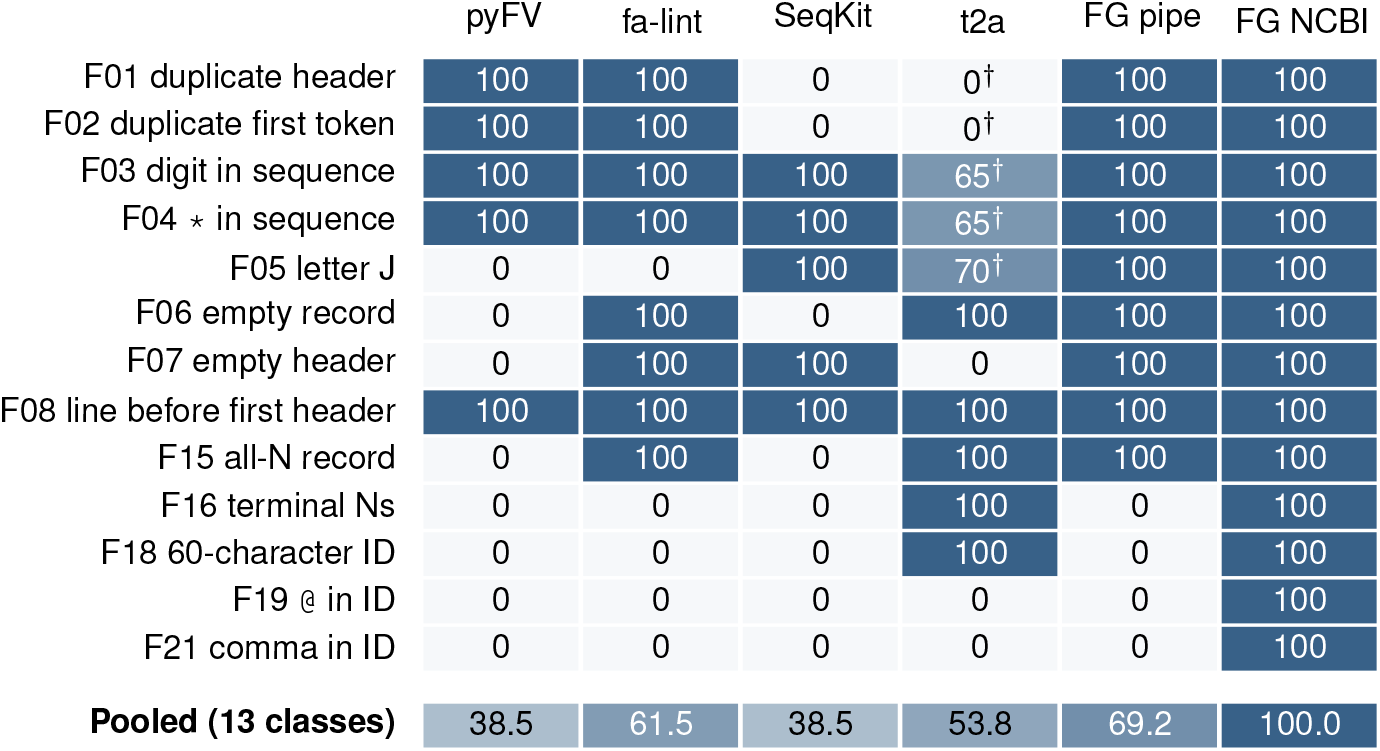
Detection of 13 classes of injected fault in real assemblies, by tool (percentage of the 40 files per class flagged; 20 backgrounds *×* 2 seeds). pyFV, py_fasta_validator; t2a, table2asn (error level); FG, FastaGuard v1.1.0 at the blocking level of its pipeline or NCBI submission gate. *†* table2asn gave no result within 60 s on some or all files of the class. The fault list was chosen by the authors and overlaps FastaGuard’s checks.

A minimal workflow pattern is therefore: run FastaGuard with `--gate pipeline` and an output bundle; inspect `gate.can_continue` in JSON (for example with `jq`); only then launch QUAST, BUSCO, BlobToolKit, CheckM, annotation, or official validators. If the gate fails, fix the FASTA-level blockers first. If the gate continues with WARN advisories, keep the findings as routing hints for which interpretive tool to prioritize next. In practice, a PASS or non-blocking WARN means only that the selected FastaGuard gate policy was satisfied and that the FASTA can be routed to appropriate downstream tools. It does not establish assembly correctness, biological completeness, contamination status, taxonomy, annotation quality, or archive acceptance. That boundary is deliberate: FastaGuard owns FASTA-level preflight and triage; interpretive tools own interpretive questions.

## 4 Discussion

FastaGuard’s contribution is a pipeline-native FASTA preflight *contract*: streaming validation, explainable findings, stable schemas, explicit gates, and packaging paths that workflow authors can adopt without reinventing per-pipeline scripts. Relative to SeqKit and SeqFu, FastaGuard trades general sequence manipulation for an opinionated QC verdict and gate policy; relative to syntax validators such as py_fasta_validator and fa-lint, it adds severity-graded, explainable findings and a structured report (Fig. 2 and Table 2). Relative to QUAST, BUSCO, BlobToolKit, and CheckM, it runs earlier and answers a different question: whether the FASTA is safe to spend those tools on. Relative to MultiQC, it is a producer of aggregatable content rather than an aggregator. Positioning the tool as preflight—rather than as a substitute for those ecosystem components—keeps it complementary to them.

The evaluation adds three points to that positioning. First, the pipeline gate’s stop rate on public assemblies is dominated by one rule: the high-N blocker stopped 109 of 247 (44.1%) of scaffold-level and none of the contig-level assemblies, and in 97 of 138 stops the per-record threshold alone was responsible. The NCBI gate’s stops are different in kind: public assemblies are survivors of repository validation, and the 14 stops are real violations of NCBI rules (short records and terminal Ns) in files the repositories hold, which we recounted without FastaGuard code. The pipeline gate’s stops are policy; laboratories that work with scaffolded genomes will want to tune or demote that rule, and whether it should block by default is a design question this survey raises and does not settle. Second, the validators largely catch different things. On injected faults, hazards of NCBI identifier and terminal-N rules (F16, F18, F19, F21) were caught by none of py_fasta_validator, fa-lint and SeqKit; table2asn reported two of them as errors, flagged a comma only as a warning and accepted @, and the pipeline gate does not block any of them by design, while the NCBI gate does. Conversely table2asn reports undeclared runs of Ns that FastaGuard reports as advisories. Third, downstream tools are neither uniformly strict nor uniformly forgiving. The defects FastaGuard blocks include all 12 silent alterations we observed (for example a space in a sequence lengthening the reported sequence by a base in three tools, or a leading byte-order mark making QUAST drop a contig), but the work the tools did before failing was small on a bacterial genome, and the one measurable saving was the reference-declaration mismatch of the `reference` mode, about a minute of mapping and sorting. The case for an early gate on these results therefore rests on explicit stop/go semantics and on the few silent alterations, not on large time savings on small genomes.

Limitations follow from that scope and from the evaluation design. FastaGuard does not discover biological contamination, does not certify archive acceptance, and does not replace official validators or FCS-style screens. The survey covers public NCBI assemblies at one date; they have already passed repository validation, assemblies above 1.5 Gb were excluded, and a stop means that a default policy fired, not that the file is unusable. The injected-fault and matrix defect lists were chosen by the authors and overlap FastaGuard’s own checks, the injected backgrounds were selected because no tool flagged them, the matrix covers one bacterial genome with sentinels that record accounting only, and the resampling intervals over backgrounds are degenerate because outcomes are fixed by class: these figures show capability on those faults, not accuracy on real submissions. Two of six pre-stated predictions about the survey were not met. All timings come from one macOS arm64 machine (and Linux containers for the gate experiment) that was not idle; absolute times should be re-measured on the target system. The adjudication of survey disagreements was done by one unblinded reviewer, and the registered second review is not done. Finding thresholds remain heuristics. The behaviour described here corresponds to release v1.1.0; later changes are recorded in the project changelog.

## Author contributions

Following CRediT, E.E.: conceptualisation, software, validation, investigation, data curation, and writing of the original draft. J.-F.M.: supervision and writing through review and editing. Both authors read and approved the final manuscript.

## AI usage disclosure

AI coding assistants (Cursor-based agents and Anthropic Claude Code) were used for planning, manuscript scaffolding, repository evidence cross-checks, prose drafting support, and revision. For the first version, the software, code base, and benchmarking results were completed first, and the manuscript was drafted solely from that code base and those results, not from a free-form prompt. For the revision, an AI assistant (Anthropic Claude Code) drafted the pre-registered analysis protocol, wrote and ran the analysis code under the first author’s direction (each data download and each analysis was approved by the first author before it ran), and drafted the new Results, Discussion and Supplementary Note from the recorded result files. The authors reviewed all output and remain responsible for the manuscript, code, evidence interpretation, and all scientific claims.

## Acknowledgements

We thank packaging and workflow maintainers who exercised Bioconda, BioContainers, nf-core module, and Snakemake wrapper integrations for FastaGuard.

## Data availability

FastaGuard source code, schemas, finding catalog, and documentation are openly available at https://github.com/ehsanestaji/FastaGuard under the MIT license (cited release tag `v1.1.0`). Committed public-evidence summaries used in Table 1 are archived under `docs/evidence/` (v0.6.0 portable JSON/TSV pack, plus the v1.0.0 re-run summary) in the repository; the rows can be regenerated with `scripts/collect_evidence.py`, which retrieves the NCBI assemblies by accession with the NCBI Datasets command-line tool and records input SHA-256 digests in the TSV. The pre-registered protocol (frozen 2026-10-27; SHA-256 in the Supplementary Note), the analysis scripts, manifests, per-assembly and per-file result tables and file hashes are available at Zenodo (doi:10.5281/zenodo.23729530). The survey assemblies are listed by accession and were retrieved from NCBI; injected files are regenerated deterministically from them; no sequence data are redistributed. No new biological samples were generated for this study; public NCBI assemblies are used under their existing data-access terms.

