## Supplementary Note for "FastaGuard: FASTA preflight QC for modern bioinformatics pipelines"

FastaGuard v1.1.0 on public assemblies, injected faults, downstream tools and workflow gates

### Supplementary Note

*Supplement to "FastaGuard: FASTA preflight QC for modern bioinformatics pipelines" (bioRxiv BIORXIV/2026/761563, revised version). DRAFT for author review: sentences in double square brackets mark facts that are not settled yet.*

#### S0 Scope and how to read this note

This note reports four analyses of FastaGuard v1.1.0 that were specified in a protocol before any data were collected: **S**, a survey of public NCBI assemblies; **F**, fault injection with ground truth; **M**, a matrix of FASTA defects against seven downstream tools; and **M2**, a replication of the gate-placement experiment with ten matched pairs per workflow engine. Every endpoint named in the protocol is reported here in whichever direction it fell, including two of six predictions about S that were wrong.

Three limits apply throughout. First, all measurements come from one platform (macOS arm64, plus Linux containers for M2) and from the tool versions in Table S1. Second, the injected-fault list in F and the defect list in M were chosen by the authors and overlap FastaGuard's own check list, so they measure capability on those faults, not accuracy on real submissions. Third, the survey describes what the default thresholds do to public assemblies; it does not measure how many of the files FastaGuard stops are unusable for any given purpose.

**Registration.** The protocol was written by an AI assistant (Claude Code) at the first author's request, frozen by the first author on 2026-10-27 at 18:02:38 UTC as a read-only copy with SHA-256 `7763c9f2546e8767e08604ab7f31ef79d6c8f6a8efcc0ce41`[illegible] and it preceded the draw of the sample and every analysis run. It has no independent timestamp from before the analyses. A copy of it was deposited at Zenodo on 2026-10-28 (doi:10.5281/zenodo.23729530), after all four analyses had run, so that date shows when the protocol became public and does not by itself show that it preceded the data. Deviations from it, with the date, the reason, and whether results had been seen, are in Section S7.

**Labels.** *Registered* means specified in the frozen protocol. *Post hoc* means added after seeing results; such items are never used to support a registered endpoint.

### S1 Software, comparators and environment

| Item | Version and identifier |
|---|---|
| FastaGuard | 1.1.0, tag `v1.1.0`, commit `71e23cf5f743039730c05d504b5c43683ab14bc3`; macOS arm64 binary SHA-256 `47973fe68aba18df031b3cd4a88ebe941198f7db3f095ce9889765065a`[illegible] Linux aarch64 binary (M2) SHA-256 `45b25bb70cf3d7e6fa5d636e4f8451b5f2923134f42b45c3fdc861fea4`[illegible] rustc 1.95.0; Bioconda `fastaguard=1.1.0`; source archived at doi:10.5281/zenodo.23728753 (all versions doi:10.5281/zenodo.23622310) |
| py_fasta_validator | 0.6; sdist SHA-256 `ca29c20e1be31ef190f923b62ddb7fccda8c1c3ba9cf699a6ccaccce10`[illegible] |
| fa-lint | 1.2.0; archive SHA-256 `6edcb792489b18bcfd03156912d05227ac05cdf657c517041c40489566`[illegible] |
| table2asn | 1.29.324 (arm64); archive SHA-256 `348b9d5f1b05065f4e5e61b6c2350e5ff044de2d6dc14765be6ac02c00`[illegible] |
| SeqKit | 2.14.0; archive SHA-256 `7b3a6a7c9be07a06c5b308d1aad769c09762b36b387690eed8b91b00d0`[illegible] |
| NCBI Datasets CLI | 18.40.0 (assembly metadata and downloads) |
| samtools, bwa, minimap2 | 1.24, 0.7.19, 2.31 (matrix); 1.16.1 and 2.24 inside the M2 images |
| BLAST+, Picard | 2.17.0, 3.5.0 |
| QUAST | 5.3.0 (osx-64 build under Rosetta; not installable on arm64) |
| BUSCO | 6.1.0, lineage `bacteria_odb12.2` (index date 2026-05-22, md5 `e21960d275e09687a86aace486f2886a`); BUSCO 6 supports only odb12 lineages |
| Snakemake, Nextflow | 9.26.1, 25.10.0 (M2 images; Docker 29.9.2) |
| Environments | conda locks `fg-downstream` SHA-256 `6bc132e385f9dd83d7577d39ad369bf6f89d500de0f398a1f0788618e0`[illegible] `fg-quast` `50c29d3713da203bde7b4bd64ed5956e05efda54775f1b874555483595`[illegible] |

*Table S1. Versions and identifiers. FastaGuard was built from* `git archive v1.1.0` *(tag tree* `a06ec5b25e526dd7e1832f1431688885b`[illegible] *not from the GitHub release asset;* `cargo test --locked --offline` *on the tagged source passed 239 tests on macOS arm64 and the same 239 on Linux aarch64. The v1.1.0 release notes list no change to analysis or exit-code contracts relative to v1.0.0; this note evaluated v1.1.0 and did not re-run v1.0.0.*

**Decision rules.** For py_fasta_validator, fa-lint, SeqKit (`seq -v -t dna`) and table2asn the rules for "flagged" and "strict" are those of `paper/comparison/run_comparison.py`, imported unchanged (SHA-256 ffb38045752b106b1a9e19cdeb1c1e33dc08f21db84b27d0d005d8571d027da0). For FastaGuard, *flagged* means verdict not PASS and *blocked* means `gate.can_continue` is false. Two configurations are used throughout, both at default thresholds: the pipeline gate (`--profile assembly --gate pipeline`) and the NCBI gate (`--profile assembly --gate submission --submission-target ncbi`). The pipeline gate blocks on duplicate identifiers, invalid characters, invalid FASTA structure and high-N content; the NCBI gate blocks on duplicate identifiers, invalid characters, invalid FASTA structure, SeqIDs that break the NCBI rule, terminal Ns and records shorter than 200 bp. A *per-record* N fraction of at least 0.20 or a *global* N fraction of at least 0.05 triggers `high_n_rate`.

**Machine.** Apple M5 Max (18 logical cores, 128 GiB), macOS 26.7.1, AC power, local SSD, warm cache. The machine was **not idle** during timed runs: other work of the first author (Docker containers, a graph database, real-time malware scanning, another coding session) was running and was not paused. The 1-minute load average after each timed survey run had median 8.3 (10th to 90th percentile 5.5 to 15.9, maximum 50.5); the timed matrix pass had mean 20.0 and maximum 55.4; the M2 runs started at a median load of 10.4. Load is recorded on every timing row. All timing statements below are conditional on it.

### S2 Analysis S: survey of 700 public assemblies

#### S2.1 Design

*Frame.* NCBI Assembly records in the RefSeq and GenBank assembly-summary listings downloaded on 2026-10-27, with `version_status` latest, a full genome representation, assembly level Complete Genome, Chromosome, Scaffold or Contig, a usable FTP path and a listed genome size of at most 1.5 Gb. *Strata and quotas*: bacteria 200, archaea 40, fungi 100, plants 100, invertebrates 100, vertebrates 100 and protozoa 60; within each stratum half from RefSeq and half from GenBank-only records (no RefSeq counterpart). *Draw*: eligible rows sorted by accession, `random.Random(20261027 + stratum index)`, the quota plus a 25% ordered reserve per cell. A reserve item could replace a sampled item only for a technical reason (404, checksum failure, wrong file type, size cap) and never because of a tool's output; no replacement was needed. All downloads (the 700 sample assemblies and the 2 anchors) matched the MD5 in NCBI's `md5checksums.txt`, and for every run FastaGuard's recorded input SHA-256 equalled the SHA-256 of the downloaded file. GRCh38.p14 (`GCF_000001405.40`) and T2T-CHM13v2.0 (`GCF_009914755.1`) were run as 2 scaling anchors and are excluded from all prevalence figures.

*Runs.* FastaGuard (both gates) on every assembly; SeqKit `stats -a`; py_fasta_validator, fa-lint, SeqKit `seq -v` and table2asn on the 401 assemblies of at most 100 Mb (120 s limit).

*Statistics.* Wilson 95% intervals for single proportions. Population-weighted figures weight each stratum by its number of eligible records and use a bootstrap interval. Assemblies of the same species or bioproject are treated as independent, which is a limitation. Unweighted figures depend on the quotas and are labelled so.

#### S2.2 Primary endpoints

| Stratum | n | Pipeline gate stops | NCBI gate stops |
|---|---|---|---|
| Bacteria | 200 | 5/200 (2.5%, 1.1–5.7) | 1/200 (0.5%, 0.1–2.8) |
| Archaea | 40 | 1/40 (2.5%, 0.4–12.9) | 0/40 (0.0%, 0.0–8.8) |
| Fungi | 100 | 18/100 (18.0%, 11.7–26.7) | 1/100 (1.0%, 0.2–5.4) |
| Plant | 100 | 30/100 (30.0%, 21.9–39.6) | 3/100 (3.0%, 1.0–8.5) |
| Invertebrate | 100 | 25/100 (25.0%, 17.5–34.3) | 1/100 (1.0%, 0.2–5.4) |
| Vertebrate | 100 | 41/100 (41.0%, 31.9–50.8) | 6/100 (6.0%, 2.8–12.5) |
| Protozoa | 60 | 18/60 (30.0%, 19.9–42.5) | 2/60 (3.3%, 0.9–11.4) |
| **All** | 700 | 138/700 (19.7%, 16.9–22.8) | 14/700 (2.0%, 1.2–3.3) |

*Table S2. Pipeline-gate and NCBI-gate stop rate by stratum (registered endpoints S-P1 and S-P2).*

Overall, the pipeline gate stopped **138/700 (19.7%, 16.9–22.8)** of the 700 assemblies unweighted and **1.9% (bootstrap 0.6–3.9)** when weighted to the population of eligible records (the weighted figure is dominated by bacteria, which are 2.5% stopped). The NCBI gate stopped **14/700 (2.0%, 1.2–3.3)** unweighted and 0.9% (bootstrap 0.03–2.5) weighted. By source, the pipeline gate stopped 75/350 (21.4%, 17.5–26.0) of RefSeq and 63/350 (18.0%, 14.3–22.4) of GenBank-only assemblies. Verdicts at the pipeline gate were PASS 41, WARN 521 and FAIL 138.

| Stratum | Source | n | PASS/WARN/FAIL | Pipeline stops | NCBI stops |
|---|---|---|---|---|---|
| Bacteria | refseq | 100 | 13/83/4 | 4/100 (4.0%, 1.6–9.8) | 0/100 (0.0%, 0.0–3.7) |
| Bacteria | genbank-only | 100 | 2/97/1 | 1/100 (1.0%, 0.2–5.4) | 1/100 (1.0%, 0.2–5.4) |
| Archaea | refseq | 20 | 12/8/0 | 0/20 (0.0%, 0.0–16.1) | 0/20 (0.0%, 0.0–16.1) |
| Archaea | genbank-only | 20 | 0/19/1 | 1/20 (5.0%, 0.9–23.6) | 0/20 (0.0%, 0.0–16.1) |
| Fungi | refseq | 50 | 9/29/12 | 12/50 (24.0%, 14.3–37.4) | 0/50 (0.0%, 0.0–7.1) |
| Fungi | genbank-only | 50 | 3/41/6 | 6/50 (12.0%, 5.6–23.8) | 1/50 (2.0%, 0.4–10.5) |
| Plant | refseq | 50 | 1/34/15 | 15/50 (30.0%, 19.1–43.8) | 2/50 (4.0%, 1.1–13.5) |
| Plant | genbank-only | 50 | 1/34/15 | 15/50 (30.0%, 19.1–43.8) | 1/50 (2.0%, 0.4–10.5) |
| Invertebrate | refseq | 50 | 0/33/17 | 17/50 (34.0%, 22.4–47.8) | 0/50 (0.0%, 0.0–7.1) |
| Invertebrate | genbank-only | 50 | 0/42/8 | 8/50 (16.0%, 8.3–28.5) | 1/50 (2.0%, 0.4–10.5) |
| Vertebrate | refseq | 50 | 0/36/14 | 14/50 (28.0%, 17.5–41.7) | 0/50 (0.0%, 0.0–7.1) |
| Vertebrate | genbank-only | 50 | 0/23/27 | 27/50 (54.0%, 40.4–67.0) | 6/50 (12.0%, 5.6–23.8) |
| Protozoa | refseq | 30 | 0/17/13 | 13/30 (43.3%, 27.4–60.8) | 2/30 (6.7%, 1.8–21.3) |
| Protozoa | genbank-only | 30 | 0/25/5 | 5/30 (16.7%, 7.3–33.6) | 0/30 (0.0%, 0.0–11.4) |

*Table S3. Stratum by source. Cells show assemblies, PASS/WARN/FAIL at the pipeline gate, and stop rates with Wilson intervals.*

**Agreement with SeqKit (S-P3).** FastaGuard's record count, total length and N50 equalled SeqKit's on 702/702 runs each (GC within 0.01 percentage points on all); there were no mismatches to investigate. FastaGuard's total length equalled the genome size in the NCBI listing for 581/702 assemblies; the remainder were not investigated (the listing's definition can differ, and listings change).

#### S2.3 What the pipeline gate stops, and why

All 138 pipeline-gate stops are the `high_n_rate` finding; no other blocking finding occurred in the sample. *Post hoc*: in 97 of them the global N fraction was below 0.05 and the per-record rule alone caused the stop (at least one record with 20% or more N), and in 41 the global fraction was also at or above 0.05. By assembly level (post hoc) the pipeline gate stopped 0/27 (0.0%, 0.0–12.5) Complete Genome, 0/239 (0.0%, 0.0–1.6) Contig, 29/187 (15.5%, 11.0–21.4) Chromosome and 109/247 (44.1%, 38.1–50.4) Scaffold assemblies.

| Assembly level | n | Pipeline gate stops | NCBI gate stops |
|---|---|---|---|
| Complete Genome | 27 | 0/27 (0.0%, 0.0–12.5) | 0/27 (0.0%, 0.0–12.5) |
| Chromosome | 187 | 29/187 (15.5%, 11.0–21.4) | 1/187 (0.5%, 0.1–3.0) |
| Scaffold | 247 | 109/247 (44.1%, 38.1–50.4) | 11/247 (4.5%, 2.5–7.8) |
| Contig | 239 | 0/239 (0.0%, 0.0–1.6) | 2/239 (0.8%, 0.2–3.0) |

*Table S4. Stop rates by assembly level (post hoc).*

The stop rate is therefore a property of the default N rule applied to scaffolded genomes, and it should be read as policy: a scaffold with one gap-rich record is stopped by default whether or not it is fit for a given downstream use. Whether the per-record rule should be a default blocker is a design question this survey raises; it is not changed in v1.1.0.

#### S2.4 Warnings and other findings

| Finding | Assemblies | % | 95% CI |
|---|---|---|---|
| length_outliers | 605/700 | 86.4 | 83.7–88.8 |
| gc_outliers | 473/700 | 67.6 | 64.0–70.9 |
| gap_runs | 291/700 | 41.6 | 38.0–45.3 |
| gap_pattern_warnings | 272/700 | 38.9 | 35.3–42.5 |
| composite_anomalies | 189/700 | 27.0 | 23.8–30.4 |
| high_n_rate | 138/700 | 19.7 | 16.9–22.8 |
| tiny_contigs | 11/700 | 1.6 | 0.9–2.8 |
| duplicate_sequences | 8/700 | 1.1 | 0.6–2.2 |
| terminal_ns | 4/700 | 0.6 | 0.2–1.5 |
| reserved_header_chars | 3/700 | 0.4 | 0.1–1.3 |
| long_headers | 1/700 | 0.1 | 0.0–0.8 |

*Table S5. Prevalence of findings at the pipeline gate (700 assemblies). Statistical outlier findings (`length_outliers`, `gc_outliers`, `composite_anomalies`) are advisory.*

WARN is not selective on public assemblies: 521 of 700 assemblies are WARN. *Post hoc*: 249 of the 521 WARN assemblies carry a gap or terminal-N finding, 270 warn only on statistical outlier findings, and 2 on other findings. Workflows should branch on `gate.can_continue`, as the manuscript recommends, and not on the verdict.

#### S2.5 NCBI-gate stops, adjudicated

The 14 NCBI-gate stops in the sample are caused by `ncbi_genome_short_contigs` (11/700) and `terminal_ns` (4/700; one assembly has both). No SeqID rule fired. For each stop (plus the GRCh38 anchor, 15 in total) the blocking counts were recomputed with stand-alone Python that does not use FastaGuard code, streaming the downloaded file and counting records shorter than 200 bases and records that start or end with N. The counts agreed with FastaGuard's in 15 of 15 cases, so all 15 are classified (a) *documented FastaGuard policy stricter than the repository's acceptance*, none (b) genuine file defect and none (c) FastaGuard defect. **The adjudication was done by one reviewer, who was not blinded; the registered independent review of 20% of cases by a second person was not done**.

#### S2.6 Comparators on the 401 assemblies of at most 100 Mb

| Tool | Level | Assemblies of at most 100 Mb |
|---|---|---|
| py_fasta_validator | flagged | 0/401 (0.0%, 0.0–0.9) |
| fa-lint | flagged | 0/401 (0.0%, 0.0–0.9) |
| SeqKit `seq -v -t dna` | flagged | 0/401 (0.0%, 0.0–0.9) |
| table2asn | flagged (error level) | 138/401 (34.4%, 29.9–39.2) |
| FastaGuard, pipeline gate | flagged (verdict not PASS) | 361/401 (90.0%, 86.7–92.6) |
| FastaGuard, pipeline gate | blocked | 40/401 (10.0%, 7.4–13.3) |
| FastaGuard, NCBI gate | blocked | 3/401 (0.7%, 0.3–2.2) |

*Table S6. Comparator flag rates (registered secondary endpoint).*

py_fasta_validator, fa-lint and SeqKit flagged none of the 401 assemblies, which is what is expected of syntax and alphabet checks on files that passed repository validation. table2asn flagged 138/401 (34.4%, 29.9–39.2), always with the single code SEQ_INST.InternalNsInSeqRaw (138), which reports a run of Ns that the submitter must declare as a gap with table2asn options. FastaGuard's NCBI gate and table2asn agreed on 2 assemblies (both flagging, for different reasons), disagreed in the other direction on 1 (FastaGuard only), and table2asn alone flagged 136. All 136 table2asn-only files carry at least one FastaGuard N or gap finding at WARN, so FastaGuard did not report them clean. The disagreement does not fit the three registered adjudication classes; it is recorded as a fourth class (d), *requirement on the submitter, not a file defect*, added after seeing the data (Section S7). The exact McNemar test was computed as registered but is not interpreted, because the disagreement is one rule and not a sample of independent events.

#### S2.7 Runtime and memory

These figures are conditional on the load stated in Section S1.

| Quantity | Value |
|---|---|
| Wall time per assembly, median / maximum | 0.54 s / 29.4 s |
| Peak RSS, median / maximum | 8.7 MiB / 628 MiB |
| GRCh38.p14 (3,298 Mbp) | 29.4 s, 10.0 MiB |
| T2T-CHM13v2.0 | 23.3 s |
| Log-log slope of wall time on compressed size (registered model 1) | 0.84 (bootstrap 0.82–0.85) |
| Same, on Mbp (descriptive) | 0.84 (0.83–0.85) |
| Wall time ratio FastaGuard / `seqkit stats -a`, median (IQR) | 2.2 (1.9–2.4) |
| Wall time ratio FastaGuard / py__fasta__validator, median | 0.92 |
| Peak RSS per 1,000 records / per Mbp (registered model 2) | 0.74 MiB / 0.0015 MiB (interval 0.0003–0.0028) |

*Table S7. Runtime and memory of the pipeline-gate run (gzip input read directly, full report bundle written).*

Median FastaGuard wall time was 0.54 s per assembly (maximum 29.4 s for GRCh38, 3,298 Mbp), median peak resident memory 8.7 MiB (maximum 628 MiB). The median ratio of FastaGuard's wall time to `seqkit stats -a` was 2.2 (interquartile range 1.9–2.4), and to py__fasta__validator 0.92; the tools do different work and the comparison is descriptive. A synthetic grid generated locally (records 10^3 to 10^6, lengths 100 to 10,000, up to 1 Gbp) separates the two sources of memory: at 1,000 bp per record, peak memory was 9.8, 18.9, 75.6 and 653.6 MiB at 10^3, 10^4, 10^5 and 10^6 records, and it did not change with total length at a fixed record count. On the 700 real assemblies the record coefficient was 0.74 MiB per 1,000 records and the per-Mbp coefficient 0.0015 MiB (interval 0.0003–0.0028); the interval excludes zero by the registered rule, but the largest genome, 3,298 Mbp, adds only about 5 MiB. The accurate wording is that memory is dominated by the number of records, with a small residual dependence on sequence length.

#### S2.8 Build cross-check

Bioconda `fastaguard=1.1.0` gave verdict, gate status, `can_continue` and finding IDs identical to the tag build on 40/40 runs (20 assemblies drawn with seed 20261027, two gates).

#### S2.9 Registered predictions against outcomes

| ID | Prediction | Observed | Outcome |
|---|---|---|---|
| P-S1 | Fewer than 5% of repository assemblies stop at the pipeline gate. | 138/700 (19.7%, 16.9–22.8) unweighted; 1.9% (bootstrap 0.6–3.9) population-weighted | **Not met** unweighted; met weighted |
| P-S2 | At least 25% return WARN, driven by gap and terminal-N findings. | WARN 521/700; 249 with a gap or terminal-N finding, 270 outlier findings only | Rate met; **driver not met** |
| P-S3 | At least 1% fail the NCBI gate, from SeqID, short-contig or terminal-N rules. | 14/700 (2.0%, 1.2–3.3); short contigs 11/700, terminal Ns 4/700, SeqID 0 | Met (SeqID rules did not fire) |
| P-S4 | Exact agreement with SeqKit on record count, total length and N50. | 702/702 for each | Met |
| P-S5 | Wall time linear in input size (slope 0.9 to 1.1). | slope 0.84 (0.82–0.85); start-up floor of about 0.1 s; loaded machine | **Not met** |

| ID | Prediction | Observed | Outcome |
|---|---|---|---|
| P-S6 | Peak memory depends on record count, not total bases. | synthetic grid: 9.8 to 653.6 MiB across 10^3 to 10^6 records, flat in total length; real data: per-Mbp interval 0.0003–0.0028 MiB | Supported, with a small residual length term |

*Table S8. Pre-stated predictions for S. P-S1 (unweighted) and P-S5 were not met; P-S2 was met on its rate but not on its stated driver.*

### S3 Analysis F: fault injection with ground truth

#### S3.1 Design

*Backgrounds.* 20 survey assemblies of at most 50 Mb were selected by agreement of all six tool configurations: an assembly was kept only if none of FastaGuard (both gates), py_fasta_validator, fa-lint, SeqKit and table2asn flagged it. The rule needed interpretation, fixed before the selector ran (Section S7): eligible assemblies (370) were shuffled with `random.Random(20261027)`; quotas were 10 bacteria (at least 3 with two or more replicons), 4 archaea and 6 fungi or protozoa; a 200-candidate budget applied to the first pass, which accepted 19 of 20; the registered relaxation to “none blocks” then supplied the remaining 1, an *Escherichia coli* complete genome that FastaGuard flags with a `length_outliers` warning but does not block. Selecting on agreement favours contig-level assemblies without long N runs, so specificity on real files is not estimated here; it comes from S.

| Accession | Group | Organism | Level | Records | Mb | Pass |
|---|---|---|---|---|---|---|
| GCF_054095835.1 | archaea | Candidatus Nitrosopumilus sp. bin_7KS | Contig | 11 | 1.71 | flagged |
| GCF_054657575.1 | archaea | Stetteria hydrogenophila | Contig | 2 | 1.36 | flagged |
| GCF_900314615.1 | archaea | uncultured Methanobrevibacter sp. | Scaffold | 57 | 2.43 | flagged |
| GCF_902384015.1 | archaea | Methanosphaera stadtmanae | Complete Genome | 1 | 1.77 | flagged |
| GCA_018630425.1 | bacteria | Staphylococcus aureus | Contig | 2 | 2.85 | flagged |
| GCA_026001605.1 | bacteria | Escherichia coli | Complete Genome | 5 | 5.42 | strict |
| GCF_000465465.2 | bacteria | Bacillus sp. NSP9.1 | Complete Genome | 1 | 4.54 | flagged |
| GCF_001010485.1 | bacteria | Dehalococcoides sp. UCH007 | Complete Genome | 1 | 1.47 | flagged |
| GCF_006370475.1 | bacteria | Escherichia coli | Contig | 2 | 4.89 | flagged |
| GCF_024029775.1 | bacteria | Fructobacillus americanaquae | Complete Genome | 1 | 1.41 | flagged |
| GCF_048972155.1 | bacteria | Klebsiella pneumoniae | Complete Genome | 3 | 5.81 | flagged |
| GCF_053348975.1 | bacteria | Klebsiella michiganensis | Complete Genome | 3 | 6.02 | flagged |
| GCF_057654775.1 | bacteria | Xylella fastidiosa subsp. fastidiosa | Complete Genome | 1 | 2.57 | flagged |

| Accession | Group | Organism | Level | Records | Mb | Pass |
|---|---|---|---|---|---|---|
| GCF_900475365.1 | bacteria | Streptococcus sobrinus | Complete Genome | 1 | 2.15 | flagged |
| GCA_030573695.1 | fungi or protozoa | Meyerozyma carpophila | Contig | 16 | 10.31 | flagged |
| GCA_034642105.1 | fungi or protozoa | Yarrowia lipolytica | Chromosome | 10 | 20.65 | flagged |
| GCF_000280035.1 | fungi or protozoa | Encephalitozoon romaleae SJ-2008 | Chromosome | 13 | 2.19 | flagged |
| GCF_001761485.1 | fungi or protozoa | Yarrowia lipolytica | Complete Genome | 6 | 20.50 | flagged |
| GCF_028827525.1 | fungi or protozoa | Penicillium daleae | Contig | 11 | 39.69 | flagged |
| GCF_028829835.1 | fungi or protozoa | Penicillium mononematosum | Contig | 6 | 30.17 | flagged |

*Table S9. The 20 backgrounds. Pass: criterion level at which the background qualified.*

*Injector.* `inject.py` takes (background, condition, seed), uses `random.Random("accession|condition|seed")` with seeds 1 and 2, and applies exactly one condition per file. The 20 backgrounds, 26 conditions and 2 seeds give **1,040 mutated files**, plus the 20 unmodified originals (all byte-identical to the downloads). Thirteen conditions are scored as errors, seven are advisory and not scored, three are a dose series of N blocks, and three are benign. The injector, manifests and FastaGuard's expected outcome per class were frozen and hashed before any tool was run on a mutated file.

| ID | Transformation | Scored as |
|---|---|---|
| F01 | append a record with the header of an existing record plus 1,000 bp from the background | error |
| F02 | append a record whose first token equals an existing record's, with a different description | error |
| F03 | replace one base with `7` | error |
| F04 | replace one base with `*` | error |
| F05 | replace one base with `J` | error |
| F06 | insert a header-only record | error |
| F07 | insert a record with the bare header `>` | error |
| F08 | insert one 100-bp sequence line before the first header | error |
| F09 | insert one blank line inside a record | advisory |
| F10 | rewrap one record at W+20 | advisory |
| F11 | convert all line endings to CRLF | advisory |
| F12 | prepend a UTF-8 BOM | advisory |
| F13 | insert one space inside a sequence line | advisory |
| F14a, b, c | overwrite a block of the longest record with N: 0.1%, 1%, 10% of total length | dose series |
| F15 | append a record of 1,000 N | error |
| F16 | overwrite the first 20 bases of the first record of at least 1,000 bp with N | error |
| F17 | append a record with a new ID and the exact sequence of the smallest record of at least 1,000 bp | advisory |
| F18 | rename a random record's ID to 60 random characters | error |

| ID | Transformation | Scored as |
|---|---|---|
| F19 | append `@` to a random record's ID | error |
| F20 | append a 150-bp record | advisory |
| F21 | insert a comma into a random record's ID | error |
| C1 | lower-case about 30% of bases in random intervals of at least 100 bp | benign |
| C2 | rewrap the whole file at width 80 (70 if W is 80) | benign |
| C3 | replace 0.01% of bases with IUPAC codes other than N | benign |

*Table S10. Conditions. "Error" classes are scored in sensitivity; "benign" classes in the false-alarm rate; advisory classes and the dose series are reported, not scored.*

*Statistics.* Seeds within a background are not independent, so the 20 backgrounds are the resampling unit: cluster bootstrap, 10,000 resamples, `random.Random(20261027)`, percentile 95% intervals. **The intervals are degenerate for most tools** (for example 38.5 to 38.5 for py_fasta_validator): each tool's outcome is fixed by the defect class and does not vary with the background, so resampling backgrounds adds no spread. The uncertainty that matters is the choice of classes, which the intervals do not cover.

#### S3.2 Sensitivity on the 13 error classes (F-P1)

| Class | pyFV | fa-lint | SeqKit | t2a flagged | t2a error | FG pipe flagged | FG pipe blocked | FG NCBI blocked |
|---|---|---|---|---|---|---|---|---|
| F01 | 100 | 100 | 0 | 0† | 0† | 100 | 100 | 100 |
| F02 | 100 | 100 | 0 | 0† | 0† | 100 | 100 | 100 |
| F03 | 100 | 100 | 100 | 65† | 65† | 100 | 100 | 100 |
| F04 | 100 | 100 | 100 | 65† | 65† | 100 | 100 | 100 |
| F05 | 0 | 0 | 100 | 70† | 70† | 100 | 100 | 100 |
| F06 | 0 | 100 | 0 | 100 | 100 | 100 | 100 | 100 |
| F07 | 0 | 100 | 100 | 0 | 0 | 100 | 100 | 100 |
| F08 | 100 | 100 | 100 | 100 | 100 | 100 | 100 | 100 |
| F15 | 0 | 100 | 0 | 100 | 100 | 100 | 100 | 100 |
| F16 | 0 | 0 | 0 | 100 | 100 | 100 | 0 | 100 |
| F18 | 0 | 0 | 0 | 100 | 100 | 5 | 0 | 100 |
| F19 | 0 | 0 | 0 | 0 | 0 | 5 | 0 | 100 |
| F21 | 0 | 0 | 0 | 100 | 0 | 5 | 0 | 100 |
| **Pooled** | **38.5** (38.5–38.5) | **61.5** (61.5–61.5) | **38.5** (38.5–38.5) | **61.5** (57.7–65.0) | **53.8** (50.0–57.3) | **78.1** (76.9–80.4) | **69.2** (69.2–69.2) | **100.0** (100.0–100.0) |

*Table S11. Percentage of the 40 files per class (20 backgrounds x 2 seeds) flagged. t2a, table2asn; FG, FastaGuard. † table2asn timed out (no result) on some or all files of the class; see below.*

Pooled over the 13 error classes (520 files): FastaGuard's **NCBI gate 520/520 (100.0%)**; FastaGuard's **pipeline gate blocked 360/520 (69.2%)** and flagged 406/520 (78.1%) (interval 76.9–80.4); table2asn flagged 320/520 (61.5%) (57.7–65.0) and reported an error-level result for 280/520 (53.8%) (50.0–57.3), with 120 runs giving no result; fa-lint 320/520 (61.5%); py_fasta_validator 200/520 (38.5%) and SeqKit 200/520 (38.5%). Of the 406 pipeline-gate flags, 40 are the advisory `terminal_ns` finding on the F16 files and 6 are files of the strict-level background, which FastaGuard flags before mutation; without that background the flagged-level figure is 76.9%. The pipeline gate blocks F01 to F08 and F15 and, by design, does not block terminal Ns, a 60-character SeqID, `@` in a SeqID or a comma in a SeqID; the NCBI gate blocks all 13.

*Paired differences in pooled sensitivity (F-P3), percentage points, FastaGuard minus comparator:* pipeline gate (blocked) minus table2asn (error level) +15.4 (+11.9 to +19.2); NCBI gate minus table2asn (error level) +46.2 (+42.7 to +50.0);

pipeline gate minus fa-lint +7.7 (+7.7 to +7.7); NCBI gate minus fa-lint +38.5 (+38.5 to +38.5); pipeline gate minus py_fasta_validator or SeqKit +30.8 (+30.8 to +30.8). The full set is Table S12.

| FastaGuard | Comparator | Difference (points) | 95% CI |
|---|---|---|---|
| FastaGuard pipeline gate (flagged) | py_fasta_validator | +39.6 | +38.5 to +41.9 |
| FastaGuard pipeline gate (flagged) | fa-lint | +16.5 | +15.4 to +18.9 |
| FastaGuard pipeline gate (flagged) | SeqKit | +39.6 | +38.5 to +41.9 |
| FastaGuard pipeline gate (flagged) | table2asn (flagged) | +16.5 | +12.3 to +21.5 |
| FastaGuard pipeline gate (blocked) | py_fasta_validator | +30.8 | +30.8 to +30.8 |
| FastaGuard pipeline gate (blocked) | fa-lint | +7.7 | +7.7 to +7.7 |
| FastaGuard pipeline gate (blocked) | SeqKit | +30.8 | +30.8 to +30.8 |
| FastaGuard pipeline gate (blocked) | table2asn (error level) | +15.4 | +11.9 to +19.2 |
| FastaGuard NCBI gate (flagged) | py_fasta_validator | +61.5 | +61.5 to +61.5 |
| FastaGuard NCBI gate (flagged) | fa-lint | +38.5 | +38.5 to +38.5 |
| FastaGuard NCBI gate (flagged) | SeqKit | +61.5 | +61.5 to +61.5 |
| FastaGuard NCBI gate (flagged) | table2asn (flagged) | +38.5 | +35.0 to +42.3 |
| FastaGuard NCBI gate (blocked) | py_fasta_validator | +61.5 | +61.5 to +61.5 |
| FastaGuard NCBI gate (blocked) | fa-lint | +38.5 | +38.5 to +38.5 |
| FastaGuard NCBI gate (blocked) | SeqKit | +61.5 | +61.5 to +61.5 |
| FastaGuard NCBI gate (blocked) | table2asn (error level) | +46.2 | +42.7 to +50.0 |

*Table S12. Paired differences, FastaGuard minus comparator, at the same level (flagged or strict), 10,000 cluster-bootstrap resamples.*

**table2asn no-results.** table2asn produced no result within the registered 60 s on 120 of 1,060 files: F01 40, F02 40, F03 14, F04 14, F05 12. The duplicate-identifier hang was known. A post hoc re-run at 300 s of 44 of these files (all F03 to F05 timeouts and the F01 and F02 files of the smallest background) timed out again in 44 of 44, so the no-results are not an artefact of the 60 s limit, and the lower table2asn sensitivities for F01 to F05 in Table S11 reflect runs that did not finish. On the files where it did finish, table2asn flagged every F03, F04 and F05 file (26, 26 and 28 files respectively).

#### S3.3 False alarms on benign changes (F-P2)

On the 120 benign files (soft-masking, rewrapping at a different width, 0.01% non-N IUPAC codes) py_fasta_validator, fa-lint, SeqKit and table2asn flagged 0/120, 0/120, 0/120 and 0/120. FastaGuard blocked 0/120 at the pipeline gate and 0/120 at the NCBI gate; at its flagged level it flagged 6/120 at both gates, all on the strict-level background whose unmodified original already carries a WARN (originals flagged at FastaGuard's flagged level: 1/20; every comparator and every blocked-level gate result on the originals: 0/20).

#### S3.4 Advisory classes and the N dose series (F-S1, F-S2)

| Class | pyFV | fa-lint | SeqKit | t2a flag | t2a error | FG pipe flag | FG pipe block | FG NCBI flag | FG NCBI block |
|---|---|---|---|---|---|---|---|---|---|
| F09 | 0/40 | 40/40 | 0/40 | 0/40 | 0/40 | 2/40 | 0/40 | 2/40 | 0/40 |
| F10 | 0/40 | 22/40 | 0/40 | 0/40 | 0/40 | 2/40 | 0/40 | 2/40 | 0/40 |
| F11 | 0/40 | 0/40 | 0/40 | 0/40 | 0/40 | 2/40 | 0/40 | 2/40 | 0/40 |
| F12 | 40/40 | 40/40 | 0/40 | 40/40 | 40/40 | 40/40 | 40/40 | 40/40 | 40/40 |
| F13 | 40/40 | 40/40 | 0/40 | 0/40 | 0/40 | 40/40 | 40/40 | 40/40 | 40/40 |
| F17 | 0/40 | 0/40 | 0/40 | 0/40 | 0/40 | 40/40 | 0/40 | 40/40 | 0/40 |
| F20 | 0/40 | 0/40 | 0/40 | 0/40 | 0/40 | 40/40 | 0/40 | 40/40 | 40/40 |
| All advisory (280) | 80/280 | 142/280 | 0/280 | 40/280 | 40/280 | 166/280 | 80/280 | 166/280 | 120/280 |
| F14a | 0/40 | 0/40 | 0/40 | 40/40 | 40/40 | 40/40 | 0/40 | 40/40 | 0/40 |
| F14b | 0/40 | 0/40 | 0/40 | 40/40 | 40/40 | 40/40 | 0/40 | 40/40 | 0/40 |
| F14c | 0/40 | 0/40 | 0/40 | 40/40 | 40/40 | 40/40 | 40/40 | 40/40 | 0/40 |

*Table S13. Files flagged out of 40 per class. The 2/40 FastaGuard flagged-level hits in F09 to F11 are the two files of the strict-level background, which FastaGuard flags before mutation.*

A blank line inside a record is flagged only by fa-lint, uneven wrapping by fa-lint on 22 of 40, and CRLF by no tool. A BOM is flagged by every tool except SeqKit, and a space in a sequence by py_fasta_validator, fa-lint and FastaGuard. An appended duplicate sequence and a 150-bp record are flagged only by FastaGuard (advisory at the pipeline gate; the 150-bp record blocks at the NCBI gate). An N block of 0.1%, 1% or 10% of the genome in the longest record is flagged by table2asn and FastaGuard in every file and by none of the three syntax tools; the pipeline gate blocks only at 10%, and the NCBI gate never, because `high_n_rate` is not an NCBI-gate blocker.

#### S3.5 FastaGuard against its own frozen expectations (F-S3)

All **2,080** gate results (1,040 files x 2 gates) matched the expectations in the protocol's Appendix A, which was written from the source before any injected file was run: **0 mismatches**. This shows the implementation does what its documentation says on these faults; it does not show that the faults are the right ones. The pipeline-gate blocking findings per class were: duplicates (`duplicate_ids`, `duplicate_first_token_ids`) for F01 and F02, `invalid_chars` for F03 to F05, `invalid_fasta_structure` for F06 to F08, `high_n_rate` for F15, and none for F16, F18, F19 and F21.

### S4 Analysis M: effect of FASTA defects on seven downstream tools

#### S4.1 Design

Thirteen defect classes, produced by the same injector, were applied to two backgrounds: **B1**, the *E. coli* K-12 MG1655 reference (NC_000913.3, 4,641,652 bp, one record), and **B2**, the same sequence split into 100 contigs. The retained `reference.fa` ends with one extra blank line; B1 is the same sequence without it, so that the clean control has no defect. The tools and their entry points were samtools `faidx`, bwa `index` and `mem` (200 error-free 100-bp reads), minimap2 `-a` (same reads), BLAST+ `makeblastdb -parse_seqids` with `blastdbcmd`, Picard `CreateSequenceDictionary`, QUAST (`--min-contig 0`, no reference; x86_64 under Rosetta because it is not installable on arm64) and BUSCO (genome mode, `bacteria_odb12.2`, offline, 1 CPU). The registered sentinel is what the tool reports about the records: names and lengths for the first five, and record count and total length for QUAST and BUSCO.

*Truth.* The literal records of a mutated file: first token of each header and the number of non-whitespace sequence characters (an offending character counts as a residue). Lines before the first header, blank lines, line terminators and a leading BOM are not records and are not counted, so for CRLF and BOM the truth is the content the file was meant to carry.

*Outcomes (assigned by script).* **FAIL**: non-zero exit, timeout or missing output. **EXACT**: exit 0 and the sentinel equals the truth. **ALTERED**: exit 0 and the sentinel differs from the truth (a record dropped, renamed or lengthened); the word is neutral and states a deviation from the file's literal content. EXACT and ALTERED carry **-Q** (quiet) or **-W** (warned): a line of output matching a registered warning pattern that does not occur in the same tool's clean runs. Mutated files were run twice (BUSCO once) and clean controls three times; all 14 clean controls were EXACT and no cell was unstable.

#### S4.2 Results

| Class | Defect | Bkg | samtools | bwa | minimap2 | BLAST+ | Picard | QUAST | BUSCO |
|---|---|---|---|---|---|---|---|---|---|
| M01 | duplicate identical header | B1 | A-W | E-Q | E-Q | FAIL | E-Q | E-Q | FAIL |
| M01 | duplicate identical header | B2 | A-W | E-Q | E-Q | FAIL | E-Q | E-Q | FAIL |
| M02 | duplicate first token | B1 | A-W | E-Q | E-Q | FAIL | E-Q | E-Q | FAIL |
| M02 | duplicate first token | B2 | A-W | E-Q | E-Q | FAIL | E-Q | E-Q | FAIL |
| M03 | digit in sequence | B1 | E-Q | E-Q | E-Q | A-W | E-Q | FAIL | E-Q |
| M03 | digit in sequence | B2 | E-Q | E-Q | E-Q | A-W | E-Q | FAIL | E-Q |
| M04 | * in sequence | B1 | E-Q | E-Q | E-Q | A-W | E-Q | FAIL | E-Q |
| M04 | * in sequence | B2 | E-Q | E-Q | E-Q | A-W | E-Q | FAIL | E-Q |
| M05 | letter J | B1 | E-Q | E-Q | E-Q | A-W | E-Q | FAIL | E-Q |
| M05 | letter J | B2 | E-Q | E-Q | E-Q | A-W | E-Q | FAIL | E-Q |
| M06 | empty record | B1 | FAIL | E-Q | E-W | A-W | E-Q | FAIL | A-Q |
| M06 | empty record | B2 | A-Q | E-Q | E-W | A-W | E-Q | FAIL | A-Q |
| M07 | empty header | B1 | E-Q | A-Q | E-W | FAIL | FAIL | FAIL | FAIL |
| M07 | empty header | B2 | E-Q | A-Q | E-W | FAIL | FAIL | FAIL | FAIL |
| M08 | blank line inside record | B1 | FAIL | E-Q | E-Q | E-Q | E-Q | E-Q | E-Q |
| M08 | blank line inside record | B2 | FAIL | E-Q | E-Q | E-Q | E-Q | E-Q | E-Q |
| M09 | CRLF | B1 | E-Q | E-Q | E-Q | E-Q | E-Q | E-Q | E-Q |
| M09 | CRLF | B2 | E-Q | E-Q | E-Q | E-Q | E-Q | E-Q | E-Q |
| M10 | UTF-8 BOM | B1 | FAIL | E-Q | E-Q | FAIL | FAIL | FAIL | FAIL |
| M10 | UTF-8 BOM | B2 | FAIL | E-Q | E-Q | FAIL | FAIL | A-Q | FAIL |
| M11 | space in sequence | B1 | FAIL | FAIL | A-Q | E-Q | A-Q | FAIL | A-Q |
| M11 | space in sequence | B2 | FAIL | FAIL | A-Q | E-Q | A-Q | FAIL | A-Q |
| M12 | 60-character SeqID | B1 | E-Q | E-Q | E-Q | FAIL | E-Q | E-Q | E-Q |
| M12 | 60-character SeqID | B2 | E-Q | E-Q | E-Q | FAIL | E-Q | E-Q | E-Q |

| Class | Defect | Bkg | samtools | bwa | minimap2 | BLAST+ | Picard | QUAST | BUSCO |
|---|---|---|---|---|---|---|---|---|---|
| M13 | comma in SeqID | B1 | E-Q | E-Q | E-Q | E-W | FAIL | E-Q | E-Q |
| M13 | comma in SeqID | B2 | E-Q | E-Q | E-Q | E-W | FAIL | E-Q | E-Q |

*Table S14. Outcome of every cell (13 classes x 2 backgrounds x 7 tools). FAIL = failed; E = EXACT; A = ALTERED; Q quiet, W warned.*

Of the 182 cells, 46 failed, 112 reproduced the records exactly (106 quietly, 6 with a warning) and 24 altered them (12 quietly, 12 with a warning). FastaGuard's pipeline gate blocked 18/26 of the mutated files (all of M01 to M07, M10 and M11), and both of its gates matched the protocol's expectations on all 26 files (mismatches: 0). In those 9 classes (126 cells) the tools failed on 40, altered the records on 24 (12 without any warning) and reproduced them exactly on 62. In the four classes the pipeline gate does not block (blank line, CRLF, 60-character SeqID, comma in SeqID) no tool altered the records (0 cells).

| Class | Defect | Pipeline gate blocks | FAIL | EXACT-Q | EXACT-W | ALTERED-Q | ALTERED-W |
|---|---|---|---|---|---|---|---|
| M01 | duplicate identical header | yes | 4 | 8 | 0 | 0 | 2 |
| M02 | duplicate first token | yes | 4 | 8 | 0 | 0 | 2 |
| M03 | digit in sequence | yes | 2 | 10 | 0 | 0 | 2 |
| M04 | * in sequence | yes | 2 | 10 | 0 | 0 | 2 |
| M05 | letter J | yes | 2 | 10 | 0 | 0 | 2 |
| M06 | empty record | yes | 3 | 4 | 2 | 3 | 2 |
| M07 | empty header | yes | 8 | 2 | 2 | 2 | 0 |
| M08 | blank line inside record | no | 2 | 12 | 0 | 0 | 0 |
| M09 | CRLF | no | 0 | 14 | 0 | 0 | 0 |
| M10 | UTF-8 BOM | yes | 9 | 4 | 0 | 1 | 0 |
| M11 | space in sequence | yes | 6 | 2 | 0 | 6 | 0 |
| M12 | 60-character SeqID | no | 2 | 12 | 0 | 0 | 0 |
| M13 | comma in SeqID | no | 2 | 10 | 2 | 0 | 0 |

*Table S15. Outcome counts per class, over 14 tool-background cells.*

**M-H1 (supported by the registered rule): quiet alteration exists.** There are 12 ALTERED-Q cells, all in classes the pipeline gate blocks: a space inside a sequence makes minimap2, Picard and BUSCO report a length one base longer than the sequence (6 cells; consistent with the space being counted as a residue); an empty record is silently dropped by BUSCO (2 cells) and by samtools on B2 (1); an empty header is renamed `(null)` by bwa (2); and a leading BOM makes QUAST on B2 drop the first contig, reporting 99 contigs and 4,595,235 bp instead of 100 and 4,641,652 (1). Several of these are minor, and the sentinels measure record accounting only; none was checked for downstream scientific consequences.

**M-H2 (supported by the registered rule): late failure exists.** 16 cells in blocked classes failed at a time of at least 0.5 times that tool's clean-run time (registered ratio 0.56–1.23). Their absolute times range from 10 ms to 1.1 s on a 4.6 Mb genome, so “late” is relative to a short run. The registered timing was taken at a mean load of 20.0; an exploratory re-timing of these cells with 10 interleaved repeats at a median load of 6.4 gave ratios 0.52–1.05 (all still at least 0.5), listed in Table S16.

| Tool | Bkg | Defect | t_fail | T_clean | Ratio | t_fail (re-timed) | T_clean (re-timed) | Ratio (re-timed) |
|---|---|---|---|---|---|---|---|---|
| BLAST+ | B1 | M01 duplicate identical header | 0.088 | 0.152 | 0.58 | 0.0873 | 0.1165 | 0.75 |
| BLAST+ | B2 | M01 duplicate identical header | 0.136 | 0.118 | 1.15 | 0.0877 | 0.1207 | 0.73 |
| BLAST+ | B1 | M02 duplicate first token | 0.106 | 0.152 | 0.7 | 0.0877 | 0.1165 | 0.75 |
| BLAST+ | B2 | M02 duplicate first token | 0.105 | 0.118 | 0.89 | 0.0858 | 0.1207 | 0.71 |
| BLAST+ | B1 | M07 empty header | 0.111 | 0.152 | 0.73 | 0.0876 | 0.1165 | 0.75 |
| BLAST+ | B2 | M07 empty header | 0.097 | 0.118 | 0.82 | 0.0869 | 0.1207 | 0.72 |
| bwa | B1 | M11 space in sequence | 1.104 | 0.899 | 1.23 | 0.7735 | 0.7397 | 1.05 |
| bwa | B2 | M11 space in sequence | 1.069 | 1.12 | 0.95 | 0.7668 | 0.7569 | 1.01 |
| Picard | B1 | M07 empty header | 0.625 | 0.549 | 1.14 | 0.4350 | 0.4419 | 0.98 |
| Picard | B2 | M07 empty header | 0.554 | 0.556 | 1.0 | 0.4131 | 0.4208 | 0.98 |
| Picard | B1 | M10 UTF-8 BOM | 0.404 | 0.549 | 0.74 | 0.3969 | 0.4419 | 0.90 |
| Picard | B2 | M10 UTF-8 BOM | 0.506 | 0.556 | 0.91 | 0.3961 | 0.4208 | 0.94 |
| samtools | B1 | M06 empty record | 0.021 | 0.027 | 0.77 | 0.0160 | 0.0180 | 0.89 |
| samtools | B2 | M10 UTF-8 BOM | 0.01 | 0.017 | 0.56 | 0.0081 | 0.0156 | 0.52 |
| samtools | B1 | M11 space in sequence | 0.024 | 0.027 | 0.91 | 0.0149 | 0.0180 | 0.83 |
| samtools | B2 | M11 space in sequence | 0.014 | 0.017 | 0.8 | 0.0101 | 0.0156 | 0.65 |

*Table S16. The M-H2 cells. Times in seconds; ratio is failure time over the clean-run time of the same tool and background.*

**Coverage.** Of 33 harmful cells (ALTERED-Q, or FAIL at a ratio of at least 0.5), 28 lie in classes the pipeline gate blocks, including all 12 quiet alterations. The other 5 are loud failures, none silent: samtools on a blank line (both backgrounds), BLAST+ on the 60-character SeqID (B2) and Picard on a comma in a SeqID (both backgrounds). CRLF was harmless in all seven tools. The comparison of tools should not be over-read: one organism, one entry point per tool, tool versions as in Table S1 (BUSCO 6.1.0 with an odb12 lineage differs from the BUSCO 5 and odb10 combination common in existing practice), and absolute times that are small.

#### S4.3 FastaGuard against its expectations on the M files

| | | Pipeline: | | NCBI: | | |
|---|---|---|---|---|---|---|
| Class | Defect | expected | observed | expected | observed | Match |
| M01 | duplicate identical header | block | FAIL (duplicate_ids,duplicate_first_token_ids) | block | FAIL (duplicate_ids,duplicate_first_token_ids) | yes |
| M02 | duplicate first token | block | FAIL (duplicate_ids,duplicate_first_token_ids) | block | FAIL (duplicate_ids,duplicate_first_token_ids) | yes |
| M03 | digit in sequence | block | FAIL (invalid_chars) | block | FAIL (invalid_chars) | yes |
| M04 | * in sequence | block | FAIL (invalid_chars) | block | FAIL (invalid_chars) | yes |
| M05 | letter J | block | FAIL (invalid_chars) | block | FAIL (invalid_chars) | yes |
| M06 | empty record | block | FAIL (invalid_fasta_structure) | block | FAIL (invalid_fasta_structure) | yes |
| M07 | empty header | block | FAIL (invalid_fasta_structure) | block | FAIL (invalid_fasta_structure) | yes |
| M08 | blank line inside record | not flagged | PASS (none) | not flagged | PASS (none) | yes |
| M09 | CRLF | not flagged | PASS (none) | not flagged | PASS (none) | yes |
| M10 | UTF-8 BOM | block | FAIL (invalid_fasta_structure) | block | FAIL (invalid_fasta_structure) | yes |
| M11 | space in sequence | block | FAIL (invalid_chars) | block | FAIL (invalid_chars) | yes |
| M12 | 60-character SeqID | no blocking finding | PASS (none) | block | FAIL (ncbi_genome_seqid) | yes |
| M13 | comma in SeqID | no blocking finding | PASS (none) | block | FAIL (ncbi_genome_seqid) | yes |

*Table S17. FastaGuard's observed gate result per class on both backgrounds against the frozen expectation.*

### S5 M2: placing the gate before mapping and sorting

#### S5.1 Design

The `fastaguard reference` gate-versus-late-validation experiment of 2026-09-02 was repeated with ten matched pairs per workflow engine and case, under the version-2 observation contract of `docs/workflow-savings-benchmark.md`, summarised by the unchanged `scripts/benchmark_workflow_savings.py`. In each pair the gated workflow runs `fastaguard reference` on a reference with a copied `.fai` whose first declared length is one base too long, and the late-validation baseline maps reads with minimap2, sorts and indexes with samtools, and only then runs the same check. Cases: a **real** case (*E. coli* NC_000913.3 and ENA run SRR27793280, 4,420,110 read pairs) and a **synthetic** case (a deterministic 5 Mbp reference and 20,000 reads of 150 bp). Engines: Snakemake 9.26.1 and Nextflow 25.10.0, in the cached Docker images of the 2026-09-02 campaign with the Linux build of FastaGuard v1.1.0 (built from the same tagged source tree; SHA-256 recorded and checked inside the container before every one of the 100 runs) bind-mounted over the image's older binary. Pairs ran back to back in alternating order on a Docker Desktop VM (18 CPUs, 46 GiB) on a busy host. Changes relative to the retained recipe: Snakemake's `--benchmark-extended`; Nextflow's raw trace (unrounded times and sizes); 10 pairs instead of 3; whole-workflow wall time recorded for both members of each pair.

#### S5.2 Results

| Engine | Case | Pairs | Gate wall (s) | Wall avoided (s) | Workflow wall avoided (s) | CPU-h avoided | Actual CPU avoided (s) | Peak RSS gate / late task |
|---|---|---|---|---|---|---|---|---|
| Snakemake | real | 10 | 0.050 | 61.9 (54.8–71.2) | 61.9 | 0.0344 | 67.6 | 2.2 MiB / 1.85 GiB |
| Nextflow | real | 10 | 0.058 | 53.6 (49.1–65.4) | 53.6 | 0.0298 | 134.3 | 15.0 MiB / 2.24 GiB |
| Snakemake | synthetic (corrected recipe) | 10 | 0.045 | 0.39 | 0.43 | 0.0002 | 0.0 | 1.0 MiB / 2.6 MiB |
| Nextflow | synthetic | 10 | 0.051 | 0.43 | 0.53 | 0.0003 | 0.6 | 15.7 MiB / 32.5 MiB |
| Snakemake | synthetic, recipe as retained (empty .fai) | 10 | 0.040 | 0.43 | 0.46 | 0.0003 | 0.0 | 1.7 MiB / 2.7 MiB |

*Table S18. Medians of ten matched pairs; the range of the per-pair task-summed wall time avoided is in brackets. Wall avoided = baseline task wall time minus gate wall time (task-summed, not elapsed workflow time); CPU-h = gross allocated CPU-hours avoided.*

In all 50 pairs the gate reported `can_continue=false` before any mapping started, and no baseline timed out. In the real case the gate cost 0.050 s (Snakemake) and 0.058 s (Nextflow) against 61.9 s and 53.6 s of mapping and sorting avoided, with a peak resident memory of 2.2 and 15.0 MiB for the gate against 1.85 GiB and 2.24 GiB for the late task. The 2026-09-02 campaign (three pairs, an earlier build) measured about 45 s avoided; the new medians are longer, which fits the busier machine and cannot be separated from it. The synthetic case avoids under half a second and shows placement and overhead, not savings.

**A defect in the synthetic Snakemake recipe.** The Snakefile baked into the cached image creates the mismatched `.fai` with `awk ... > fixture/reference.fa.fai` over the very file it reads, which leaves it empty (checked: 0 bytes). The gate then reports `reference_malformed_declaration` in addition to the declaration mismatch, so that recipe does not test the intended one-base mismatch. The Nextflow recipe writes a different file name and is unaffected. The synthetic case in Table S18 therefore uses a corrected Snakemake recipe (the mismatch is written to `fixture/reference.mismatch.fai`); the recipe as retained was also run for ten pairs and avoided 0.43 s. The Snakemake synthetic row published earlier in the project documentation came from this recipe; its raw files are not retained, so whether that run had the empty `.fai` cannot be checked. This was found only by looking at the gate output of one smoke run.

The contract limits what may be said: *the measured reference mismatch was stopped before mapping and sorting, avoiding the reported median work on this runner and fixture.* It is not a claim about other workflows, inputs or platforms.

### S6 Limitations (collected)

1. **Survey.** The sample is of public NCBI assemblies at a single date and is stratified, not proportional. Repository assemblies have passed validation, so low defect rates were expected. Assemblies over 1.5 Gb are excluded, so large mammalian and plant genomes are under-represented. Related assemblies are treated as independent. The unweighted pipeline-gate stop rate depends on the quotas, and the population-weighted rate is dominated by bacteria.

2. **Policy, not correctness.** A pipeline-gate stop means the default policy was triggered; it does not mean the file is unusable. No tool in this note measures how often a stopped assembly would have damaged a downstream analysis.
3. **Predictions.** Two of six predictions about the survey were wrong (Table S8). The N rule's reach was not anticipated.
4. **Timing.** All timings were taken on a busy machine, with warm caches, on one platform; absolute times should be re-measured before citing.
5. **Adjudication.** Single reviewer, not blinded; the registered second review is not done; a fourth class was added after seeing the data.
6. **Fault injection.** The fault list is author-chosen and overlaps FastaGuard's checks; backgrounds were selected for being unflagged by all tools, so they under-represent files with long N runs; intervals are degenerate and do not cover the choice of classes; table2asn did not finish on 120 files.
7. **Downstream matrix.** One organism of 4.6 Mb, seven tools at the versions listed, record-accounting sentinels only, no assessment of scientific consequences, absolute times that are small, and no cell-level predictions were registered.
8. **Gate placement.** One engine pair per case on Docker Desktop with a busy host, one fixture family, and a recipe correction (Section S5).
9. **Not covered.** Protein inputs, `compare` mode, CheckM and BlobToolKit, Linux and Windows host runs, real repository submissions and user studies.

### S7 Deviations from the registered protocol

| Date | Item | Deviation or interpretation | Results seen before? |
|---|---|---|---|
| 2026-10-27 | Timing conditions (S) | The timed survey and the synthetic grid ran on a machine that was not idle; load recorded per run. | No |
| 2026-10-28 | Typology of adjudication (S) | Class (d), requirement on the submitter, added for the 136 table2asn-only disagreements. | Yes |
| 2026-10-28 | Second review (S) | The independent review of 20% of adjudications was not done. | n/a |
| 2026-10-28 | Post hoc breakdowns (S) | Which `high_n_rate` trigger fired; what causes WARN; stop rate by assembly level. | Yes |
| 2026-10-28 | Background selection (F) | Interpretation of quotas, replicon definition and the 200-candidate budget fixed before the selector ran; the strict-level relaxation supplied one background. | Unmodified assemblies only |
| 2026-10-28 | Injector (F, M) | Open points of the injector table fixed in its docstring; truth excludes BOM and CR. | No |
| 2026-10-28 | Background B1 (M) | The retained reference's trailing blank line removed from the clean control. | Calibration only |
| 2026-10-28 | Sentinels (M) | BUSCO summary parsing corrected before freezing; BUSCO 6.1.0 run end to end on the clean control. | Clean control only |

| Date | Item | Deviation or interpretation | Results seen before? |
|---|---|---|---|
| 2026-10-28 | Warning inspection (M) | One regex miss (BLAST+ comma message): two cells changed from EXACT-Q to EXACT-W. | Yes |
| 2026-10-28 | Timing conditions (M) | Timed matrix pass on a busy machine; exploratory 10-repeat re-timing of the M-H2 cells. | Re-timing after the H2 result |
| 2026-10-28 | table2asn timeouts (F) | 120 runs hit the 60 s limit; registered handling kept; post hoc 300 s re-run of 44 files. | Yes |
| 2026-10-28 | Analysis script timing (F) | The F analysis script was written and hashed while collection was running; only a 6-row smoke test and the last five log rows had been seen. | Marginally |
| 2026-10-28 | Linux binary for M2 | The registered hash equality across operating systems is impossible; replaced by a build from the same tagged tree (46 crates fetched, 8.1 MB, with the first author's approval); hash checked in all 100 runs. | n/a |
| 2026-10-28 | M2 recipe | Benchmark-extended and raw traces; 10 pairs; workflow wall time; the synthetic Snakemake recipe corrected (empty `.fai`) with the recipe as retained also run. | A one-pair smoke run |

### S8 Reproducibility and data

All scripts, manifests, per-file and per-assembly result tables and hashes are in the project evidence record deposited at Zenodo, doi:10.5281/zenodo.23729530 (published 2026-10-28; archive `fastaguard-v1.1.0-evaluation-evidence.zip`, SHA-256 `05874f5f01c261484d790562f0caefa0ef23f53c7b422dc1e641d4b459af35e5`). No sequence data are redistributed: assemblies are retrieved by accession (the list is `survey/sample_final.tsv`), the injected files are regenerated deterministically by `faultinj/inject.py` from the downloaded backgrounds, and the matrix inputs by `matrix/build_inputs.py` from the retained E. coli reference. Table S19 lists the frozen scripts and the principal result files with their SHA-256 digests.

| File (under `paper/revision/`) | SHA-256 (first 16 hex digits) |
|---|---|
| `PROTOCOL.v1.0-frozen.md` | `7763c9f2546e8767` |
| `survey/draw_sample.py` | `7db7684f3e3cd26a` |
| `survey/sample_frozen.tsv` | `94b27787a9431c6a` |
| `survey/run_survey.py` | `dde47fdd535ecc9f` |
| `survey/analyze_survey.py` | `0045a276f84b17ca` |
| `survey/adjudicate_ncbi.py` | `e07958af6af03691` |
| `survey/adjudicate_t2a.py` | `0ee225bb26cc4544` |
| `survey/results/fg_runs.tsv` | `beaa02c2cb476ea1` |
| `survey/results/timing.tsv` | `fb42912c5633e98c` |

| File (under `paper/revision/`) | SHA-256 (first 16 hex digits) |
|---|---|
| `survey/results/comparators.tsv` | 5a68255abb007fb0 |
| `survey/results/s_summary.json` | 4d872251fb192753 |
| `faultinj/select_backgrounds.py` | b4610d7e0790d781 |
| `faultinj/inject.py` | e302b960549efaaa |
| `faultinj/build_all.py` | fd21093a4f1ad736 |
| `faultinj/manifest.jsonl` | 0fd63d483a4f3ddc |
| `faultinj/run_faultinj.py` | 86cd25dd42d176cc |
| `faultinj/analyze_faultinj.py` | 88e69750f03e4a97 |
| `faultinj/results/f_runs.tsv` | 5724d4a0f5426a48 |
| `faultinj/results/f_summary.json` | 500b3d2a614c48a2 |
| `faultinj/results/f_t2a_retry.tsv` | 2556ccf012996b1d |
| `matrix/build_inputs.py` | 850925bda211a5db |
| `matrix/tools.py` | 9b3e4fd84d3f666d |
| `matrix/run_matrix.py` | 03f3380722c4cf4d |
| `matrix/analyze_matrix.py` | ca2a4966b8eaac89 |
| `matrix/manifest.jsonl` | 5880eab905b03e5d |
| `matrix/finalize_matrix.py` | 22a3821893fd8185 |
| `matrix/retime_h2.py` | c011512bf53bb782 |
| `matrix/results/m_runs.tsv` | 355557d0752b10e2 |
| `matrix/results/m_summary.json` | 2a1f90add0918ffc |
| `matrix/results/m_cells_final.tsv` | 050c9dc0c64e6c35 |
| `m2/run_m2.py` | d97f8f923f71f4e0 |
| `m2/results/m2_pairs.tsv` | 61225d0936e23dbd |
| `m2/results/snakemake-real-summary.json` | b240a8318e7a3e69 |
| `m2/results/nextflow-real-summary.json` | 0bd4452e9a4368ce |
| `m2/results/snakemake-synthetic-summary.json` | 70d23eeab3e77e0f |
| `m2/results/nextflow-synthetic-summary.json` | 37e002c54566554d |
| `m2/results/snakemake-synthetic_asretained-summary.json` | bd4fcc20eca35644 |

*Table S19. Frozen scripts and result files.*

### S9 Use of AI assistance in this work

The protocol, the analysis scripts, the execution of the analyses and the first draft of this note were produced by an AI assistant (Anthropic Claude Code) working under the first author's direction, in a session that included approval of each download and each analysis by the first author.
